# Reduced planning depth and stronger probability discounting in alcohol and tobacco use disorder

**DOI:** 10.64898/2026.09.09.750151

**Authors:** Johannes Steffen, Sarah Schwöbel, Stefan J. Kiebel, Michael N. Smolka

## Abstract

**Background:** Substance use disorders (SUDs) are associated with impaired goal-directed decision-making, but the specific cognitive mechanisms underlying these deficits remain unclear. Forward planning, a key component of goal-directed control, requires strategies to contain computational demand and ensure feasibility through heuristics such as reducing planning depth, pruning unlikely options, and discounting probabilistic outcomes. This study investigated whether individuals with alcohol use disorder (AUD) and tobacco use disorder (TUD) differ from healthy controls in their use of these heuristics during sequential decision-making.

**Methods:** We examined participants from the general population meeting criteria for at least moderate AUD (n = 45), TUD (n = 49), or both (n = 45), and compared them with matched healthy control groups. To assess forward planning, we used a previously published probabilistic three-step Markov decision task. Computational modeling was used to infer individual planning depths and compare model variants incorporating low-probability pruning, probability discounting, or both.

**Results:** A combined model including both low-probability pruning and probability discounting best explained participants’ behavior and was subsequently used for analysis. Compared to matched healthy controls, all groups with SUD exhibited significantly reduced planning depth. Moreover, individuals with SUD tended to show stronger probability discounting, with this effect reaching statistical significance in the comorbid group.

**Conclusion:** Individuals with AUD and/or TUD engaged in forward planning using similar strategies as healthy controls but exhibited reduced planning depth and a tendency toward stronger probability discounting. These findings suggest the presence of altered planning processes in SUD, which could contribute to maladaptive decision-making. Future research is needed to clarify whether such differences reflect pre-existing cognitive traits, consequences of substance use, or both, and to explore their potential relevance for intervention.

## INTRODUCTION

Substance use is a major global health problem with tobacco and alcohol being the deadliest substances accounting for almost 15% and 5% of all deaths respectively (World Health Organization, 2024a). About 21% of the world’s population aged 15 years and older regularly smokes tobacco while 7% live with alcohol use disorder (AUD; World Health Organization, 2024a, 2024b).

These numbers signal a need for action. However, the limited understanding of fundamental cognitive mechanisms underlying deteriorating substance use trajectories is one major problem hampering the development of innovative prevention and intervention approaches (Scheier, 2015). Initially, substance use trajectories start out as goal-directed decisions to use a substance, which over many repetitions can transition into habitual substance use (Lally et al., 2010), a process proposed to underly addiction (Everitt & Robbins, 2016). A central question is why certain individuals repeatedly make such goal-directed decisions that ultimately foster the formation of habitual consumption patterns (Redish et al., 2008). We hypothesize that distinct strategic approaches during goal-directed decision-making can be identified, reflecting cognitive mechanisms that bias substance-related choices. In the present cross-sectional study, we sought to examine these strategic differences and provide deeper insight into goal-directed decision-making through computational modelling, while controlling for basic cognitive abilities known to be broadly affected through neurotoxicity of substance exposure (Brust, 2010; Durazzo et al., 2010; Glass et al., 2009).

Goal-directed decision scenarios are usually of sequential nature, i.e. goal pursuit occurs over a series of decisions where initial decisions affect the conditions of later decisions (Mattar & Lengyel, 2022). These sequential decision scenarios require forward planning in order make adaptive decisions and this process can become highly complex (Geffner, 2013; Simon & Daw, 2011). In recent decades, cognitive neuroscience has advanced our understanding of how humans manage such decision complexity, in part through computational frameworks derived from model-based reinforcement learning (Sutton & Barto, 2018; for a review see Mattar & Lengyel, 2022), which formalize processes such as forward planning. Here, forward planning is described as an internal forward simulation through the decision tree of possible sequences of actions and their anticipated consequences to find the sequence yielding the most preferred outcome (Sutton & Barto, 2018). In most real-world scenarios however, humans do not have the cognitive- and time resources to plan through the whole decision tree (Callaway et al., 2022; Snider et al., 2015). Therefore, they are expected to ignore parts of the information and use efficient approximations, so-called heuristics (Gigerenzer & Gaissmaier, 2011). In forward planning, a dominant source of complexity is the planning depth as the decision tree grows exponentially with deeper planning (Callaway et al., 2022; Geffner, 2013). In our previous study, we showed using a newly developed planning task that humans limit their planning depth (Steffen et al., 2023) but found no difference in participants with mostly mild-to-moderate AUD (Steffen et al., 2025). In the present study, we focus on moderate-to-severe cases and extend our approach by considering an additional planning heuristic, i.e. pruning of low-probability outcomes. Previous studies have shown that humans use such pruning strategies to reduce planning complexity (Huys et al., 2012, 2015). We therefore tested whether individuals with SUD differ from healthy controls (HC) in their use of both depth limitation and pruning during forward planning.

A further bias that has been demonstrated in humans is probability discounting. Previous research has shown that human decision-makers discount the overall value of an outcome proportional to the odds against receiving it (Green & Myerson, 2004) which has also been demonstrated during forward planning (Sass et al., 2025). In the context of SUD, this bias has only been investigated with non-sequential choice tasks and most studies indicate stronger probability discounting in SUD groups compared to healthy controls (for an overview see Garami & Moustafa, 2020; Harrison et al., 2018). Stronger probability discounting and low-probability pruning could explain the preference for the reliable effects of substance use and the neglect for rare but severe negative consequences in addiction.

In a prior study investigating healthy participants playing our planning task, we confirmed that they most likely used all of the afore-mentioned heuristics (Sass et al., 2025). As all of these heuristics have explanatory potential for aspects of goal-directed decision-making in SUD, we therefore investigated whether participants with SUD also exhibit these strategies and if there may be differences compared to HC participants. We are not aware of any previous study examining heuristics during forward planning in SUD. To identify these heuristics and infer parameters like planning depth limit or probability discounting rate, we used a refined version of our planning task and applied computational modelling (Montague et al., 2012). To determine whether differences in forward planning strategies represent a general phenomenon of SUD, we assessed samples of participants from the general population fulfilling at least moderate AUD, tobacco use disorder (TUD) or both as well as matched healthy controls. We particularly aimed at testing the following three hypotheses:

1. Participants with AUD and/or TUD apply the following heuristics during forward planning: limitation of planning depth, low-probability pruning and probability discounting
2. The planning depth limit is further reduced in participants with AUD and/or TUD
3. Probability discounting is more pronounced in participants with AUD and/or TUD

## MATERIAL AND METHODS

### Participants and Procedure

We recruited potential participants from the general population in three ways: by recontacting the basic cohort of the German Collaborative Research Centre 265 (Heinz et al., 2020), via online advertisements in large German cities and by systematically contacting addiction counseling centers in Germany. Interested individuals first underwent an online pre-screening filtering for age and risky use of illegal drugs and were offered time slots to choose from for the telephone screening. On appointment date, after giving informed online consent, participants were screened for the following exclusion criteria: age outside the range of 18-65 years, insufficient language-, visual or motor skills for the study, current intake of any psychotropic drugs or current treatment of a severe psychiatric disorder. Included participants were subsequently diagnosed for the fulfilment of the criteria of AUD and TUD during the past 12 months according to the fifth edition of the Diagnostic and Statistical Manual of Mental Disorders (DSM-5; American Psychiatric Association, 2013) with a structured clinical interview (SCID-5-CV; First, 2016). For the three SUD groups, participants had to fulfill at least a moderate AUD- and/or mild TUD diagnosis while HC participants were only included when currently not fulfilling any SUD criterion. Finally, the main part of the study was carried out by the participants on their private computers at home as a browser-based online experiment within 4 weeks following the telephone screening. The online experiment consisted of the main planning task, a neuropsychological test battery with three cognitive tasks as well as sociodemographic and psychological questionnaires. Overall, the procedure took 2.5-3 hours (0.5-1 hours telephone screening and 2 hours online experiment) and participants received 25-35 € compensation depending on their performance in the planning task. The study was approved by the ethics committee (EK536122019) of the Dresden University of Technology and performed in accordance with relevant guidelines and regulations.

To reduce selection bias, we recruited a greater sample of HC participants over a wide range of values for sociodemographic variables for later one-to-one propensity score matching to each of the clinical groups (see below). Before analyzing the complete datasets, we excluded participants whose performance was below expected results for random behavior in any of the tasks (see Table S1 for more information). This led to the exclusion of seven HC datasets, three datasets of the AUD group, two datasets of the TUD group and one dataset of the comorbid AUD and TUD group. In total, we included *n* = 111 complete datasets for the HC group, *n* = 45 for the AUD group, *n* = 49 for the TUD group and *n* = 45 for the AUD and TUD group.

### Propensity Score Matching

Prior to further analyses, propensity score matching was performed separately for each of the SUD groups. This procedure minimized group differences of four sociodemographic variables (age, gender, higher education and working status) by finding matched dataset pairs with similar values. For each SUD dataset, a best-matching HC 5 dataset was selected. We used the PsmPy package (Kline & Luo, 2022) version 0.3.13, which computed logistic propensity scores from the sociodemographic variables as a joint distance measure for the matching. Because matching was conducted separately for each clinical group, the same HC participant could theoretically be selected as a match in more than one comparison. Out of the *n* = 111 complete HC datasets, *n* = 72 were finally used as matches. Propensity score matching successfully reduced group differences of all clinical groups in nearly all sociodemographic variables to small effect sizes (below 0.2; see Figure S1) and none of these variables showed a significant difference between groups (see Table 1).

**Table 1.**
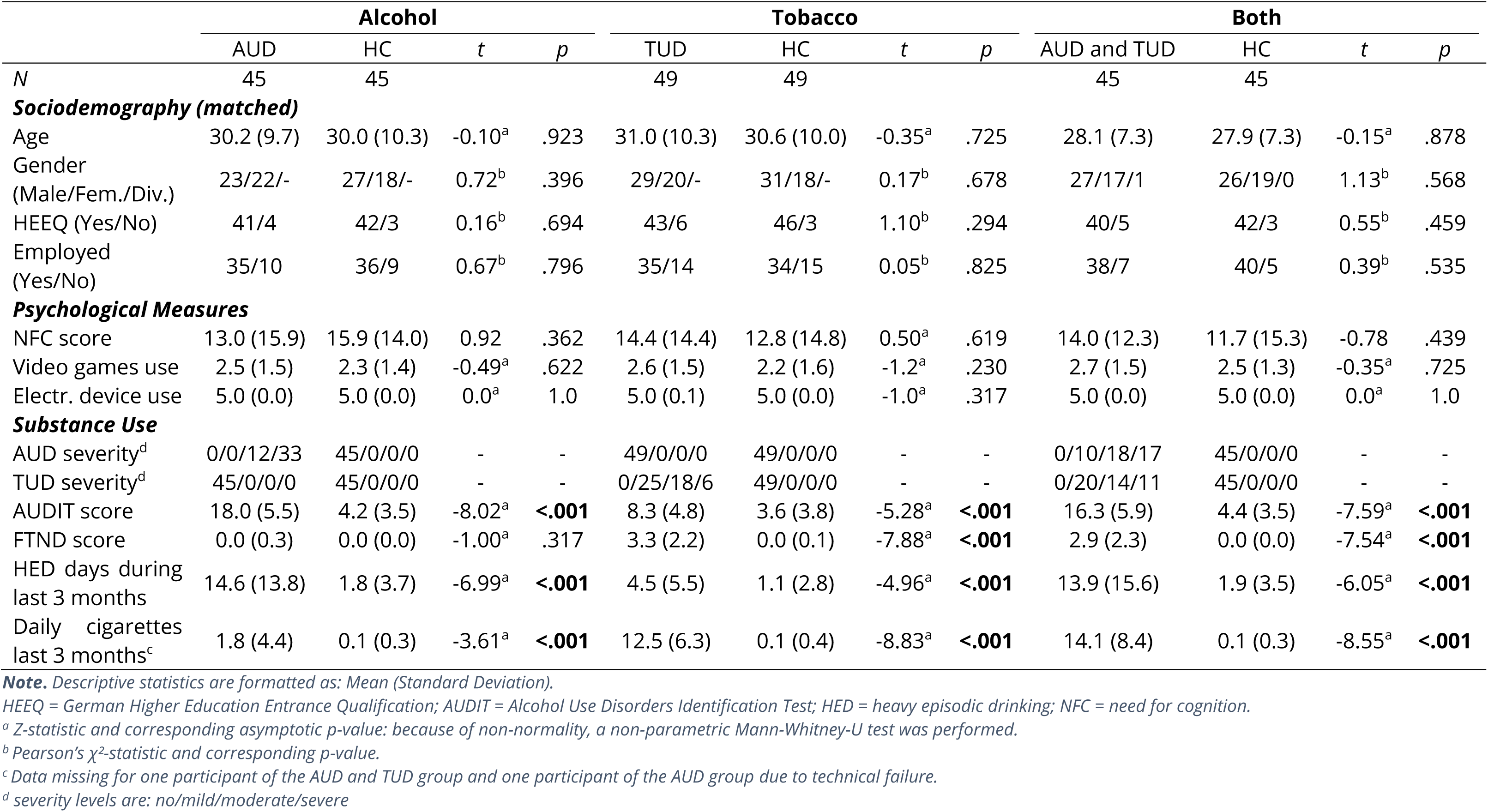
Sample Characteristics of the three SUD groups and matched healthy controls.

### Psychological Measures and Cognitive Tasks

To evaluate participants’ disposition toward effortful thinking as a potential covariate of forward planning, we employed the 16-item German version of the Need of Cognition Scale (Bless et al., 1994). For all SUD groups, there was no significant difference in NFC scores compared to their respective matched HC sample. Moreover, we asked participants for their frequency of playing computer games or using electronic devices as another potential confounder for task performance. Similarly, there were no significant group differences between SUD- and matched HC groups (see Table 1).

Lastly, we controlled for the following three basic cognitive abilities involved in forward planning: processing speed, spatial working memory and logical reasoning. We used a picture matching task (Identical Pictures Task, IDP; Lindenberger et al., 1993) for processing speed, a Spatial Working Memory Task (SWM; Nagel et al., 2008) and the 12-item form of Raven’s Advanced Progressive Matrices (RAV; Bors & Stokes, 1998) for logical reasoning. For a detailed description of these three cognitive tasks, please refer to (Steffen et al., 2023). Performances in these tasks are reported in the results section (also see Table 1) and included in the statistical analysis of forward planning outcomes.

### Probabilistic Planning Task

The Space Adventure Task (SAT) is a probabilistic multi-step decision-making task that requires forward planning of action sequences in order to maximize reward (Steffen et al., 2023). In each trial (referred to as a mini-block), participants navigated a spaceship through a planetary system by visiting a sequence of three planets (**Figure 1** A). Each planetary system consisted of six planets, selected from five distinct types, each associated with specific fuel point gains or losses (**Figure 1** B). The primary objective was to accumulate the highest possible amount of fuel across 140 unique mini-blocks, each with varying planet configurations and starting positions. A fuel bar displayed at the top of the screen tracked cumulative fuel across trials. Within each mini-block, participants were given three action steps to maneuver the spaceship, with the number of remaining steps indicated centrally on the screen with green rectangles.

**Figure 1.**
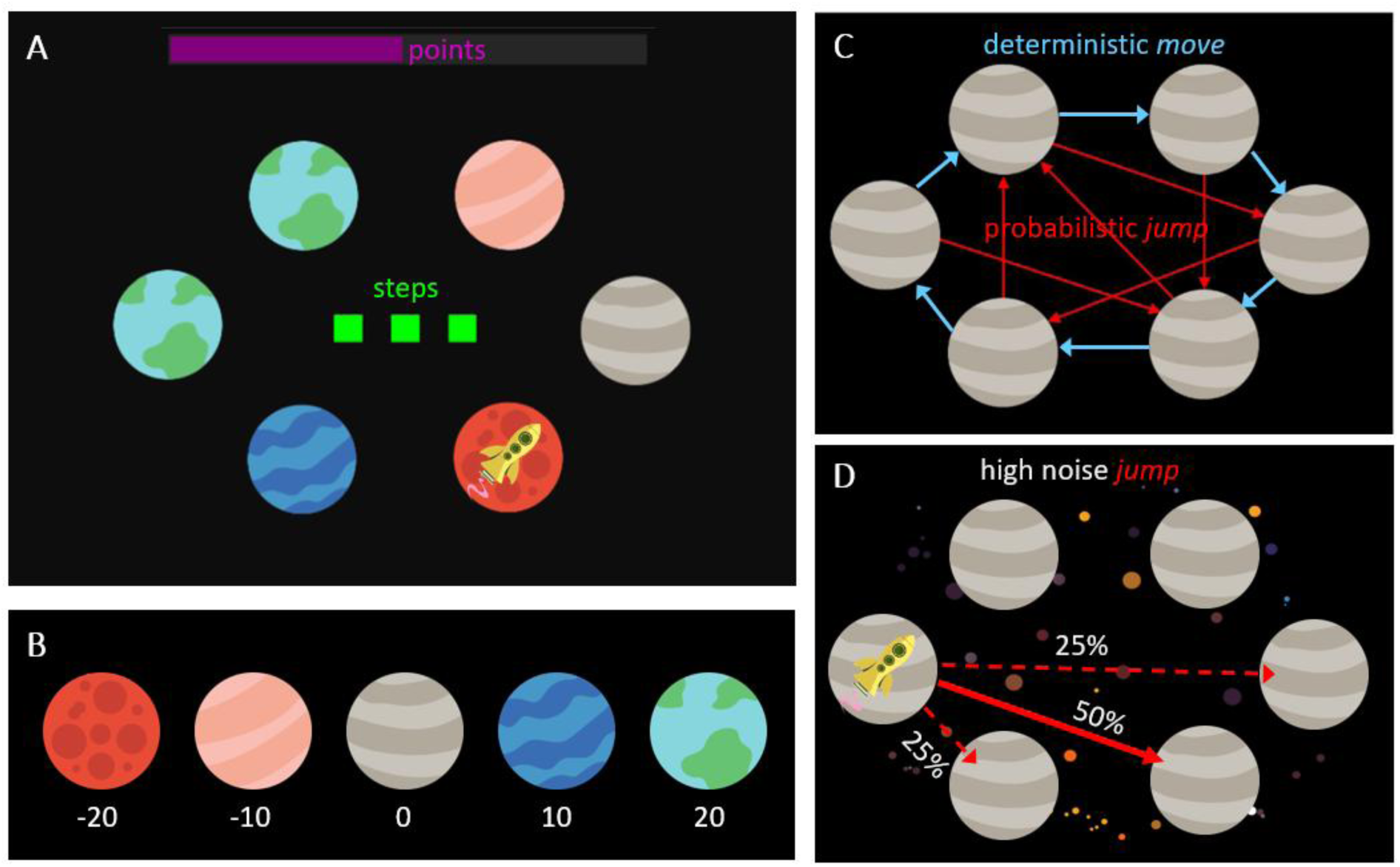
Schematic of the Space Adventure Task (adapted from Steffen et al., 2023). **(A)** Example mini-block with three action steps (green squares) under low-noise conditions (black background). The yellow rocket indicates the current location while the fuel bar at the top depicts total points accumulated so far throughout the task. **(B)** The five planet types with their respective reward values. **(C)** Transition rules for move (clockwise blue arrows) and jump (red arrow pattern) indicating the target from each possible start position. **(D)** Visualization of the probabilistic jump transition with three possible outcomes (target or one of the neighboring planets). The example shows high-noise conditions (asteroids in background) with a probability of 50% to land on the target and 25% to land on each of the neighboring planets. Under low-noise conditions, the probability to land on the target was 90% and 5% for each of the neighboring planets.

At each decision point, participants chose between two types of actions: a deterministic “move” to the adjacent planet in a clockwise direction, or a probabilistic “jump” to a non-adjacent planet based on a predefined transition structure (**Figure 1** C). The “move” action had a fixed transition probability of 100%, while the “jump” action involved uncertainty: there was a high-probability transition to the target planet (either 90% or 50%, depending on condition), and two lower-probability transitions (either 5% or 25%, respectively) to one of the two neighboring planets of the target (**Figure 1** D). These neighboring outcomes were equally likely.

Transition uncertainty was manipulated across two noise conditions. In the low-noise condition, the jump action led to the target planet with 90% probability and to each neighboring planet with 5% probability. In the high-noise condition, the target transition occurred with only 50% probability, and each neighboring planet was visited with 25% probability. The current noise condition was visually cued by the presence (high-noise) or absence (low-noise) of asteroid icons in the background (**Figure 1** D). Importantly, transition probabilities were governed solely by the noise condition and remained constant across all states. Noise conditions alternated pseudo-randomly every 3–6 mini-blocks, resulting in 70 high-noise and 70 low-noise mini-blocks across the experiment.

Participants were instructed to identify the optimal action sequence in each mini-block to maximize fuel gain. Action choices and planning times (i.e., the latency between stimulus onset and selection of the first action) were recorded for each mini-block.

Prior to the main experiment, participants underwent structured training to familiarize themselves with the task goal, the structure of the transition matrix, and the characteristics of the noise conditions. While the probabilistic nature of the jump action was explained, the exact transition probabilities were not disclosed explicitly; instead, participants were expected to infer them from experience during training. Training included sample mini-blocks with performance feedback to support learning of the task mechanics.

The task was controlled via a standard computer keyboard. The deterministic move and probabilistic jump actions were executed using the ‘Y’ and ‘M’ keys on a German QWERTZ keyboard, respectively, pressed with the left and right index fingers. The assignment of keys to actions was counterbalanced across participants to control for motor biases.

### Cognitive Models: Overview

To investigate potential planning mechanisms employed by participants, we implemented and fit four different reinforcement learning (RL) models to behavioral data based on our previous publication (Sass et al., 2025). Each model formalizes a distinct hypothesized strategy or bias that may govern participants’ choices during the task: full planning, discounted full planning, low-probability pruning, and discounted low-probability pruning. These models were evaluated using the Widely Applicable Information Criterion (WAIC) to determine which strategy best accounted for the observed behavior. Across all models, we infer each participant’s effective planning depth, i.e. the average number of steps ahead in the decision tree that their strategy implies they considered.

We begin by outlining the theoretical assumptions that distinguish the four planning models (see **Figure 2**). A detailed formal description of their mathematical implementation follows in the subsequent section. The models differ in several key respects: the extent of the decision tree that is explored at each step, whether transition probabilities are considered, and how probabilistic outcomes are subjectively evaluated by the decision-maker.

**Figure 2.**
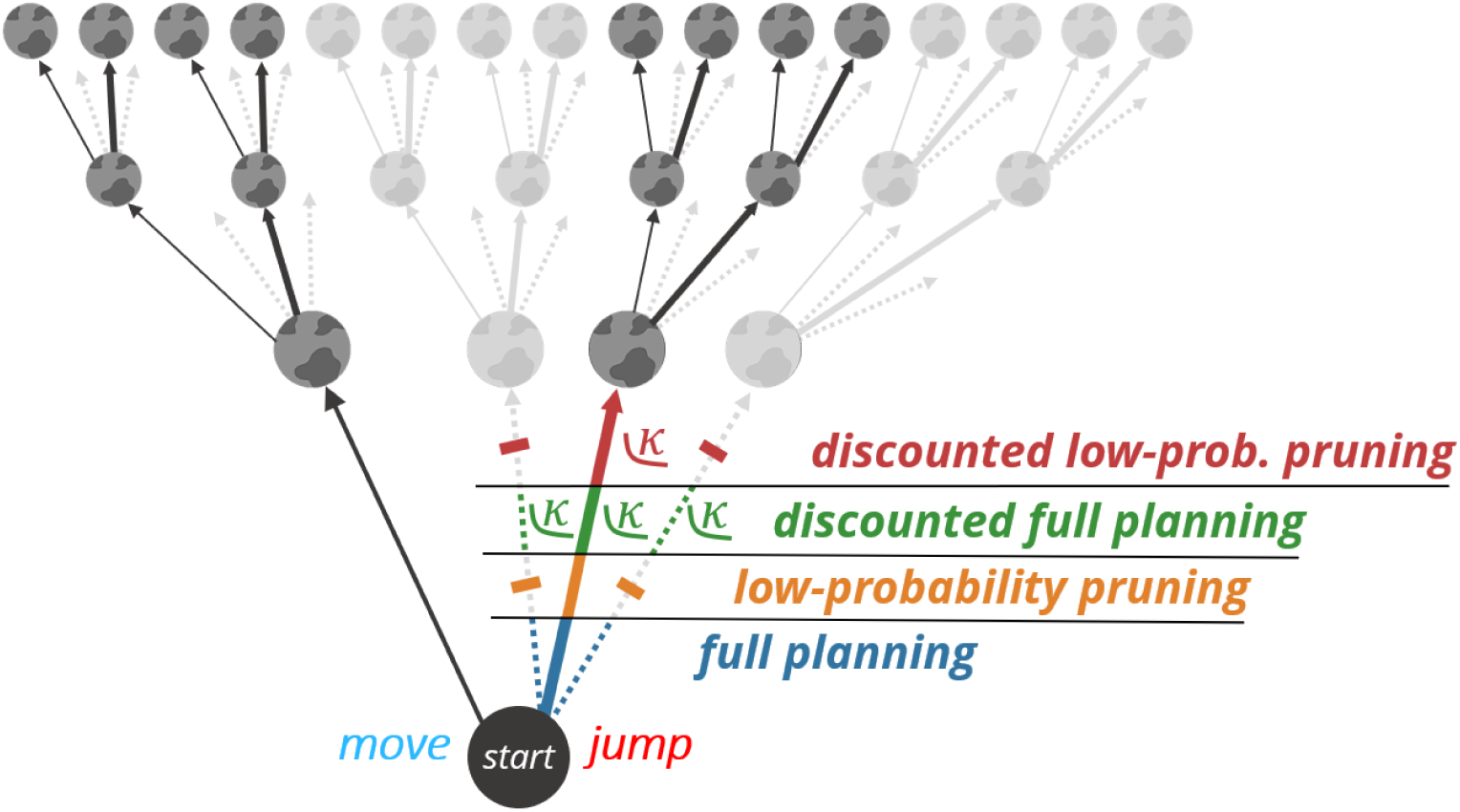
Visualization of the four forward planning strategies in the decision tree of the SAT. In each mini-block, participants could plan 3 steps ahead and choose between a deterministic move and a probabilistic jump action. The jump action led to the main target with a probability of p=0.9 or 0.5 (thick arrow) and to neighboring planets with a probability of p=0.05 or 0.25 each (dotted arrows). The full planning strategy (blue) evaluated all the branches of the tree with their accurate probabilities. The low-probability pruning strategy (orange) disregarded the low-probability transitions to the neighboring planets in all the levels of the decision tree (greyed out) and treated the transition to the main target as deterministic. The discounted full planning strategy (green) evaluated all branches but applied hyperbolic probability discounting to the jump transitions. Discounting was applied globally to the overall Q-value of the jump action with a participant-specific discounting parameter (κ). This corresponds to discounting the jump transitions of the first step in the decision tree. The discounted low-probability pruning strategy (red) combined probability discounting with pruning of the low-probability transitions. Thus, discounting was effectively only applied to the main transition of the first step in the decision tree with a participant-specific discounting parameter (κ).

The full planning model assumes that participants evaluate all possible future states at each step of the decision tree, incorporating accurate transition probabilities. In contrast, the low-probability pruning model proposes a computational heuristic that minimizes cognitive load by ignoring low-probability jump outcomes and treating high-probability transitions as effectively deterministic. This heuristic reduces the branching factor at each step, thereby simplifying the state-space representation significantly.

We also examined biased variants of these two strategies by incorporating probability discounting mechanisms. In these models, subjective values are assigned to probabilistic outcomes using a hyperbolic discounting function (Rachlin et al., 1991), reflecting individual tendencies to devalue uncertain rewards. In the discounted full planning model, all possible jump outcomes are considered, but the overall value of the jump action is hyperbolically discounted based on the probability of the main transition. In the discounted low-probability pruning model, only the high-probability jump transition is retained and subjected to probability discounting, while low-probability branches are pruned from the decision tree. Together, these models distinguish between limitations of cognitive effort during planning and biases in how participants evaluate uncertainty.

### Cognitive Models: Mathematical Formalization

The alternative cognitive models are based on the default full planning model, which was adapted from the model introduced by Steffen et al. (2023). As participants were not informed about the exact transition probabilities of the jump action, we assumed they might continuously update their belief about these transitions with temporal difference learning. However, in all prior studies using our task, parameter fitting yielded very small values for the corresponding learning rate parameter alpha (Steffen et al., 2023, 2025). This suggests that participants might have already well approximated the correct probabilities during training. Removing this learning mechanism did not reduce model likelihood or change WAIC scores (see Figure S3). Therefore, we adopted the more parsimonious model for all subsequent analyses. In the following, we now describe the mathematical formalization of the four models.

**Full planning:** In the full planning model, participants’ action choices per mini-block were formalized with a mixture model of three model-based RL agents with planning depth 1, 2 and 3, respectively. Each agent was implemented with a fully informed model of the environment, including the position of the rocket, the available actions a ∈ {′*move’*, ‘*jump*’} and states *S* (the planet configuration of the mini-block), the current noise condition with its transition probabilities *P(S_t+1_|S_t_, a_t_)* for reaching a subsequent state *S_t+1_* from a given state *s_t_* with action *a_t_*, as well as the immediate reward of reaching each state *r(s)* indicated by the planet types of the current configuration.

To decide which action *a* to choose in a specific state *s*, the agents planned through all the trajectories of the decision tree within the limit of their respective planning depth *d* and computed the expected cumulative reward each action would yield at the end of the mini-block known as Q-values, *Q(a_t_, s_t_, d)*. Mathematically, this optimal planning procedure was implemented with the value iteration algorithm (Sutton & Barto, 2018).

Finally, participants’ choices were modelled probabilistically with a softmax function, based on the computed *Q*-values for the two actions: the higher the relative value of one of the actions, the higher the probability of selecting that action. For our case of two available actions, this corresponded to a sigmoid transformation *σ(x)* of the difference between the *Q*-values, Δ*Q(s_t_, d)*. Choice probabilities were thus defined as:

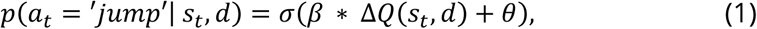

where

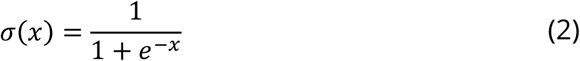

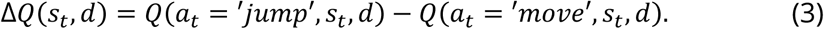

Here, we included two established parameters that further modify choice probabilities. First, the inverse response noise *β* which controlled the extent to which differences in *Q*-values affected action selection. If *β = 0,* actions are selected with a constant probability independent of outcomes, while higher values of *β* represented higher probability to select the action with the highest *Q*-value. Second, the parameter *θ* which denoted an a priori response bias, where positive values imply a bias towards choosing ‘jump’. Hence, the full planning model contained three free parameters: *β*, *θ* and *d*.

**Discounted full planning:** this model extends the full planning model by probability discounting, i.e. a discounting of all outcomes of the probabilistic jump action. Discounting was applied at the root of the decision tree (see Figure 2), so all action sequences starting with a jump were discounted according to the probability of the jump transition. Therefore, this model differs from the full planning model in the computation of the relative *Q*-values:

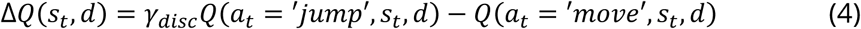

Here, the expected cumulative reward of the jump action is discounted by a discounting factor *γ_disc_* which follows a typical hyperbolic discounting function (Green & Myerson, 2004):

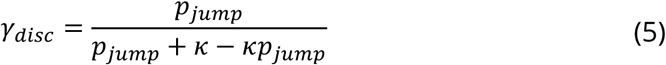

where *p_j__ump_* denotes the probability of the high-probability jump transition *p(s_t+1_* = *‘target’*|*s_t_,a_t_ = ‘jump‘)* and *κ* denotes the individual discounting parameter. In the case of *κ* = 0, there was no discounting. Higher values of *κ* led to stronger probability discounting and, hence, to lower subjective values of all outcomes of the probabilistic jump action. The *κ*-values were limited at a maximum of 30, as values beyond this threshold did not offer additional information. The discounted full planning model contained four free parameters: *β*, *θ, κ* and *d*.

**Low-probability pruning:** this model modifies the full planning model by pruning all branches of low-probability transitions of jump actions in the whole decision tree and treating jump actions as deterministic. Mathematically, this corresponds to a modification of the agents’ belief about the transition probabilities of the jump action, such that *p(s_t+1_ ^= ‘^target’*|*S_t_,a_t_* = *‘jump’)* = 1. By changing the transition matrix, this subsequently also affects the computation of *Q*-values. The low-probability pruning model contained three free parameters: *β, θ* and *d*.

**Discounted low-probability pruning:** this model modifies the full planning model by introducing both, probability discounting of jump outcomes and pruning of low-probability branches. Jump transitions were again treated as deterministic but action sequences starting with a jump are additionally discounted depending on the true transition probabilities: *P_ju__mp_ = 0.9* for low noise and *P_j__ump_ =* 0.5 for high noise. Mathematically, *Q*-values are calculated as in the low-probability pruning planning model and relative *Q*-values are computed as in the discounted full planning model (see Eq. (4)). The discounted low-probability pruning model contained four free parameters: *β*, *θ, κ* and *d*.

### Planning Depth and Parameter Inference

To infer from participants’ choices the posterior distributions over the described free model parameters, we extended the behavioral models described above by a hierarchical Bayesian model over the free parameters using the same hierarchical extension for each of the four behavioral models. As an analytical solution for the posteriors of the free parameters is intractable, we conducted stochastic variational inference using Pyro v1.5.2 (Bingham et al., 2019) to approximate the posterior distributions. As described in the previous sections, the behavioral models contained two or three parameters on the subject-level (inverse response noise *β*, response bias *θ* and discounting rate *κ* if applicable) and planning depth *d* on the mini-block level. The inference procedure can be imagined as a two-step procedure. Firstly, approximate posteriors of *β*, *θ* and *κ* were computed for each of the three RL agent models with planning depth *d ∈ {1,2,3}*. Secondly, these agents with their respective inferred parameter distributions for inverse response noise *β*, response bias *θ* and discounting rate *κ* were used to infer the posterior over planning depth *d*. For this purpose, the response likelihoods of the three agents with their respective planning depth were aggregated in a mixture model being weighted by the probability each planning depth has in mini-block *b*. This mixture model is expressed as:

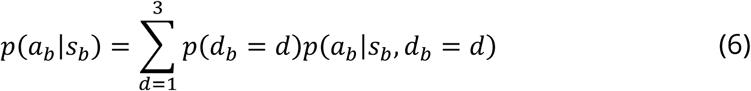

The response likelihoods based on the mixture model were maximized during inference. Importantly, we assumed that forward planning should mostly happen before the first action of each mini-block. Hence for inference, we constrained behavioral data only to the first choice in each mini-block and therefore focused on the planning depth before the first action. We placed a uniform Dirichlet prior over planning depths. As a result, the posterior samples of planning depth *d* for each mini-block form a categorical distribution. We computed the expected value of this distribution to obtain the mean planning depth *d̄* for each mini-block. In the following, we refer to this parameter simply as planning depth. For more details on the computational model and inference procedure see Steffen et al. (2022).

### Model Comparison

To assess model fit, we computed the Widely Applicable Information Criterion (WAIC; Watanabe & Opper, 2010):

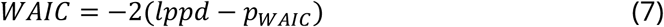

This criterion estimates the expected log pointwise predictive density (*lppd*) and includes a penalty term for overfitting risk (*p_WAIC_*), which is related to the variance of the log pointwise predictive density. WAIC therefore inherently balances fit and complexity of Bayesian models (Gelman et al., 2014).

### Statistical Analyses

All statistical analyses were conducted using IBM SPSS Statistics (version 28). We performed separate analyses for each of the SUD groups, comparing participants with AUD, TUD and both with their respective matched HC group. The distribution of continuous variables, including planning time, reaction time (RT) and performance in the tasks, estimated model parameters as well as sociodemographic and substance use characteristics, was assessed for each group using the Shapiro–Wilk test to evaluate normality, and Levene’s test to assess homogeneity of variance between groups. In cases where the assumption of normality was violated for at least one of the groups, non-parametric group comparisons were carried out using the Mann-Whitney U test. When heterogeneity of variance was detected, Welch’s t-test was employed to compare groups. In the absence of these violations, two-tailed independent-samples t-tests were used. For categorical variables, group differences were assessed using Pearson’s chi-squared (*χ*²) test. A significance level of 0.05 was applied for all statistical tests. For the case of more than one significant group effect in the model parameters, we performed post-hoc pairwise comparisons using Z test for two independent effect size estimates. The effect of the noise condition in the SAT was analyzed with linear mixed effects models with random intercept and random slopes and a condition-by-group interaction term. To analyze the 14 relationship between planning depth, planning time and performances in cognitive tasks, we conducted multiple linear regression analyses of planning depths with group indicator, IDP-, SWM- and RAV task performance as well as planning time as predictors.

In addition to the separate analyses for each of the SUD groups, we also investigated the relationship of the SUD conditions. Specifically, we tested if the effects of the AUD and TUD condition are more than additive in the comorbid case. For this goal, we performed factorial analyses of SAT performance, planning time, planning depth *d* and the model parameters (*β*, *θ, κ)* regressing each of these outcomes on indicator variables for the presence of AUD, TUD and an interaction term (see Table S3) for all included participants.

Post-hoc, we analyzed the association of SUD severities with planning depth in all SUD patients with a linear regression using the sums of fulfilled diagnostic criteria for AUD and TUD respectively as well as an interaction term as predictors (see Table S4). To further explore the influence of consumption levels, we also performed a linear regression of planning depth with numbers of smoked cigarettes and heavy episodic drinking days during the last month as well as an interaction term as predictors (see Table S5).

## RESULTS

We begin by comparing each of the SUD groups with their respective matched HC group in the behavioral outcomes of the Space Adventure Task and the cognitive tasks. We then continue with reporting the model comparison and analyses of the model parameters of the winning model. Descriptive statistics and group comparisons are presented in Table 2.

**Table 2.**
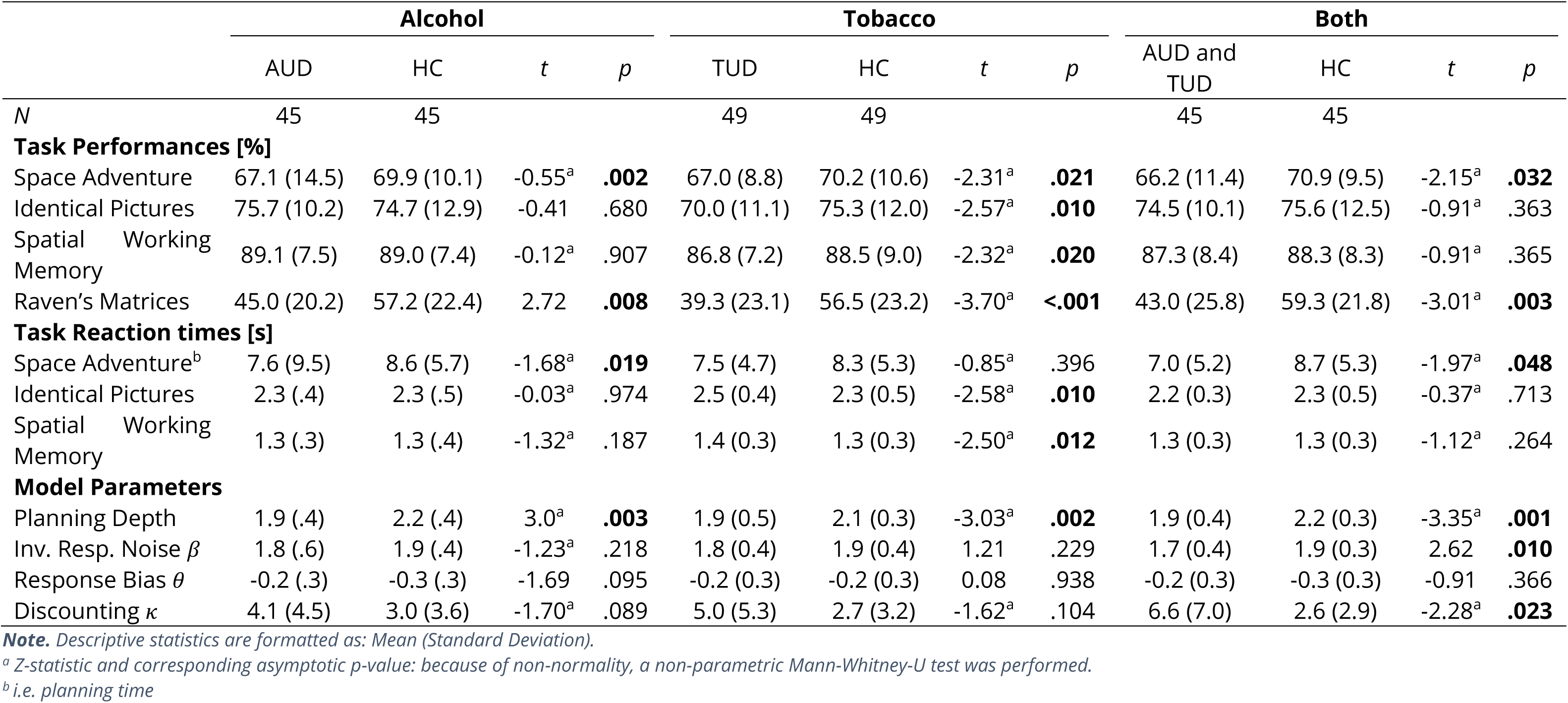
Descriptive Statistics and group comparison of task outcomes and model parameters.

### Behavioral Outcomes

We found lower performances in the planning task for all subgroups with substance use disorder, i.e. AUD (*Z* = -0.55, *p* = .002), TUD (*Z* = -2.31, *p* = .021) as well as AUD and TUD (*Z* = -2.15, *p* = .032), compared to their matched HC group. Subgroups with alcohol problems also showed significantly shorter planning times than HC participants, i.e. AUD (*Z* = -1.68, *p* = .019) as well as AUD and TUD (*Z* = -1.97, *p* = .048). For logical reasoning assessed by the raven’s task, performances were decreased in all SUD groups, i.e. AUD (*t*(88) = 2.72, *p* = .008), TUD (*Z* = -3.70, *p* < .001) as well as AUD and TUD (*Z* = -3.01, *p* = .003). Additionally, the TUD group achieved lower performances in the spatial working memory task (*Z* = -2.32, *p* = .020) and in the identical pictures task indicating lower processing speed (*Z* = -2.57, *p* = .010) while taking on average more time to respond in these two tasks (SWM: *Z* = -2.50, *p* = .012; IDP: *Z* = -2.58, *p* = .010).

### Model Comparison of Planning Strategies

The comparison of the four different planning strategy models revealed that the combined model with pruning of low-probability branches as well as probability discounting fit the data best (see Figure 3). All further analyses of model parameters are therefore based on this winning model. Separate model comparison for each SUD group and the pooled group of all HC participants included through matching showed similar results for each group (see Figure S4). Only for the AUD group, the fit of the discounted low-probability pruning model was indistinguishable from the full planning and discounted full planning model, so there was no single best fitting model.

**Figure 3.**
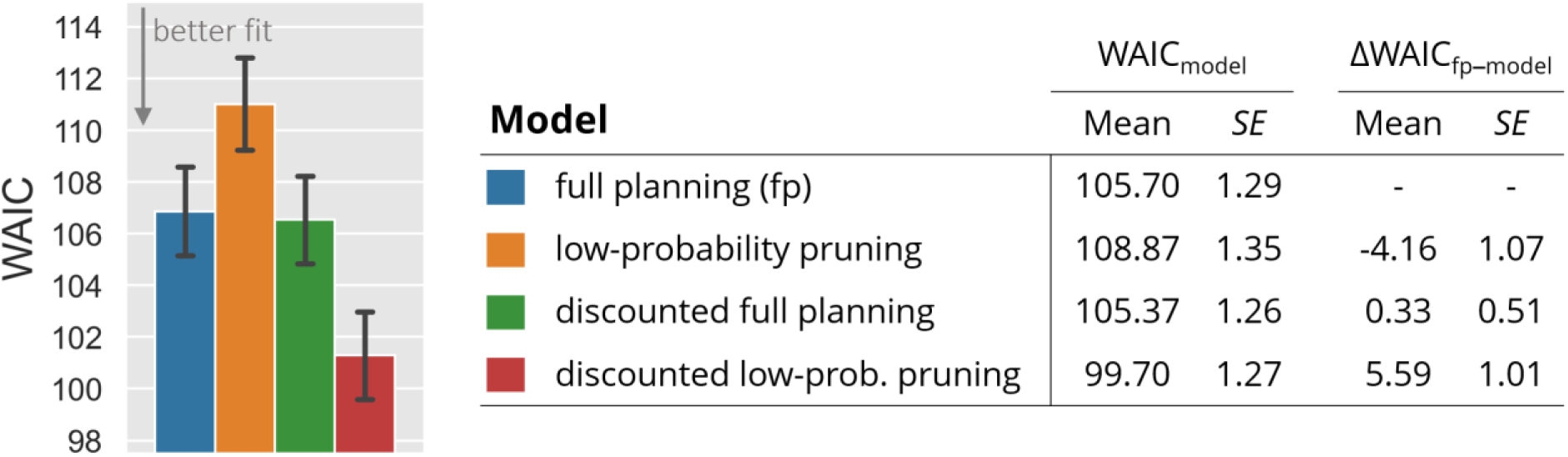
Model comparison of the four planning strategies with the Widely Applicable Information Criterion (WAIC), across SUD participants and HC participants included during matching. The full planning strategy (fp, blue) evaluated all the branches of the decision tree with their accurate probabilities. The low-probability pruning strategy (orange) disregarded the low-probability transitions to the neighboring planets. The discounted full planning strategy (green) evaluated all branches but applied hyperbolic probability discounting to the jump transitions and the discounted low-probability pruning strategy (red) combined probability discounting with pruning of the low-probability transitions. Error bars indicate the standard error of the mean of participants’ values (SE). The barplot’s values are also depicted in the WAICmodel column table. The last column of the table presents the mean pairwise WAIC differences of the participants’ values between the null model (full planning) and each of the other models alongside the corresponding standard error of the mean (SE).

### Planning Depth and model parameters

The distribution and group comparison of all model parameters is illustrated in Figure 4. The planning depth parameter was reduced in all subgroups with SUD compared to their matched HC group, i.e. for AUD (*Z* = 3.00, *p* = .003), TUD (*Z* = -3.03, *p* = .002) as well as AUD and TUD (*Z* = -3.35, *p* = .001). Post-hoc pairwise Z-tests revealed that there was no significant difference between any of these three effects (see Table S2). Furthermore, we found that the comorbid group with AUD and TUD also showed higher response noise as indicated by the parameter *β* (*Z* = 2.62, *p* = .010) and stronger probability discounting as indicated by the discounting parameter *κ* (*Z* = -2.28, *p* = .023) as the HC participants. Across all participants, there was a strong positive correlation of performance in the planning task with planning depth (*r* = 0.63, *p* < .001) as well as with inverse response noise *β* (*r* = 0.72, *p* < .001) and a strong negative correlation with the discounting parameter *κ* (*r* = -0.60, *p* < .001).

**Figure 4.**
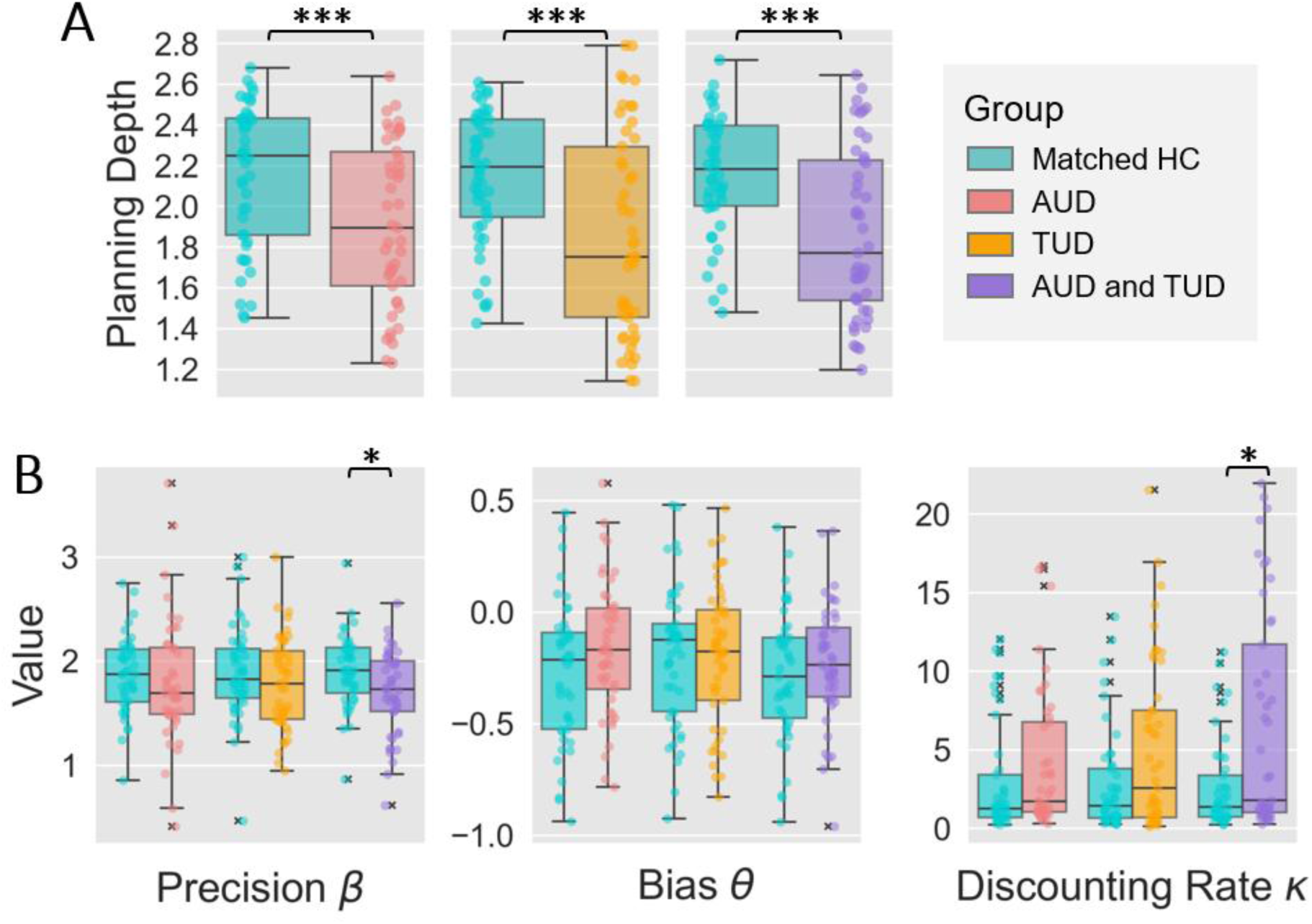
Boxplots of inferred model parameter values per subject comparing the groups with AUD/TUD/both to their respective matched healthy control (HC) group. **(A)** Averaged planning depths per subject. **(B)** Additional model parameters precision β, bias θ and probability discounting rate κ per subject.

To analyze the effect of the noise condition on planning depth, we performed separate linear mixed effects model analyses for each SUD group also showing reduced planning depths in SUD compared to matched control participants. Moreover, the analyses revealed that across groups participants, under high noise, marginally reduced their planning depth on average by 0.06. This noise effect as well as overall mean of planning depths represented by the intercept showed considerable variation among participants as indicated by the variance parameter *η*. For detailed statistics, please refer to Table 3.

**Table 3.**
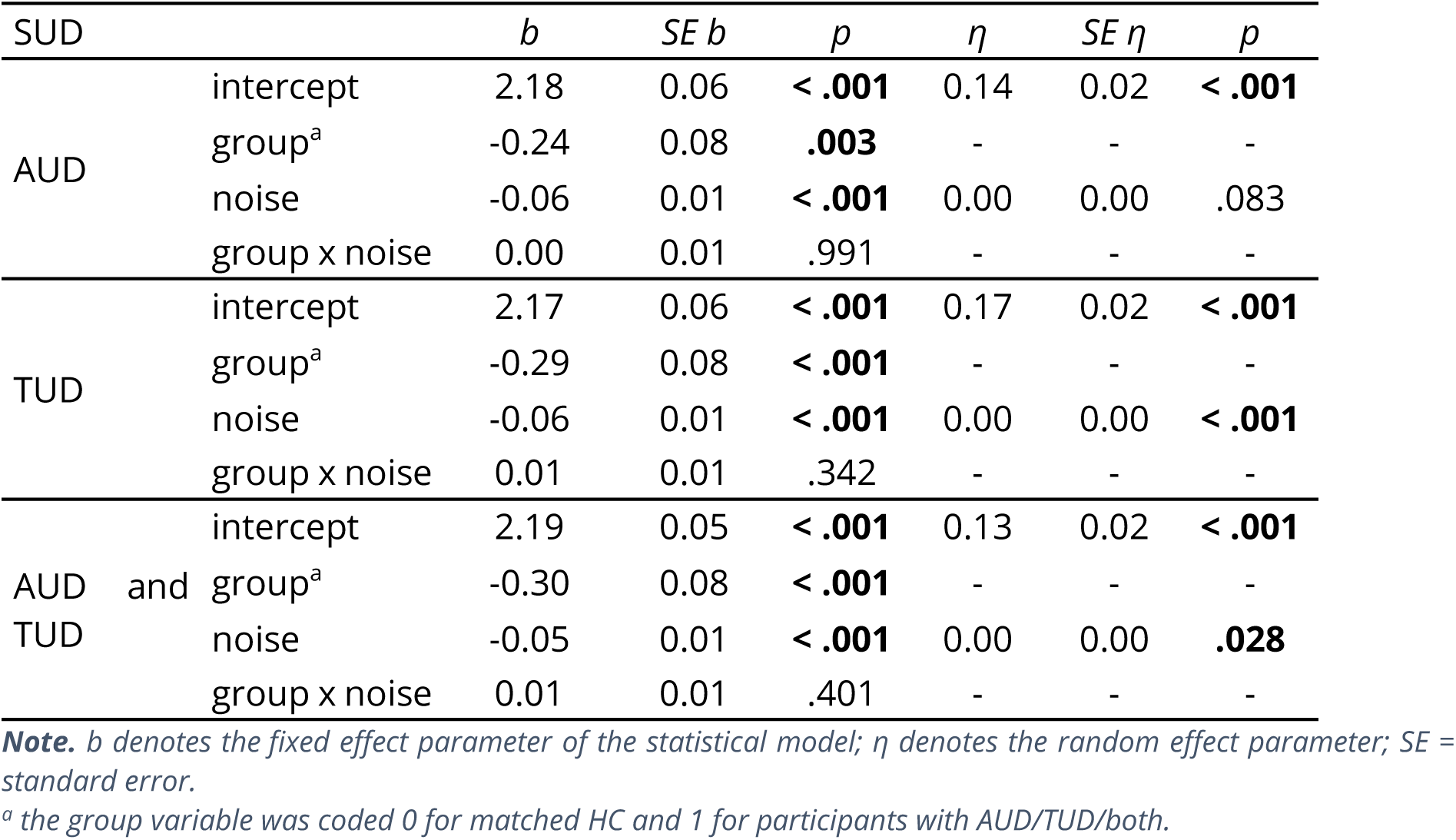
Estimates of Linear Mixed Effects Model for Planning Depth.

To investigate the relationship of planning depth with planning time and the cognitive tasks, we performed a multiple linear regression for each SUD group and relied on non-parametric bootstrapping (BCa; Efron & Tibshirani, 1994) because of non-normal distributions of planning depths (see Table 4). For all SUD groups, we found a significant positive association of planning time and planning depth, i.e. for the AUD (*p* = .038, BCa 95% CI [0.01, 0.07]), TUD (*p* < .001, BCa 95% CI [0.02, 0.07]) and the AUD and TUD group (*p* < .001, BCa 95% CI [0.02, 0.06]). Despite the inclusion of the additional predictors, the group effect of planning depth was still significant for the AUD group (*p* = .015, BCa 95% CI [-0.36, 0.0]) and the AUD and TUD group (*p* = .006, BCa 95% CI [-0.32, -0.06]). In the TUD group, we found a significant positive association with the logical reasoning performance in the raven‘s task (*p* = .001, BCa 95% CI [0.0, 0.01]) and here, the group effect got borderline significant when accounting for planning time and performances in cognitive tasks (*p* = .059, BCa 95% CI [-0.28, 0.0]). Across all participants, there were positive correlations of planning depth with planning time (*r* = 0.52, *p* < .001) as well as with logical reasoning (*r* = 0.38, *p* < .001; see FIGURE S5 for a full correlation table).

**Table 4.**
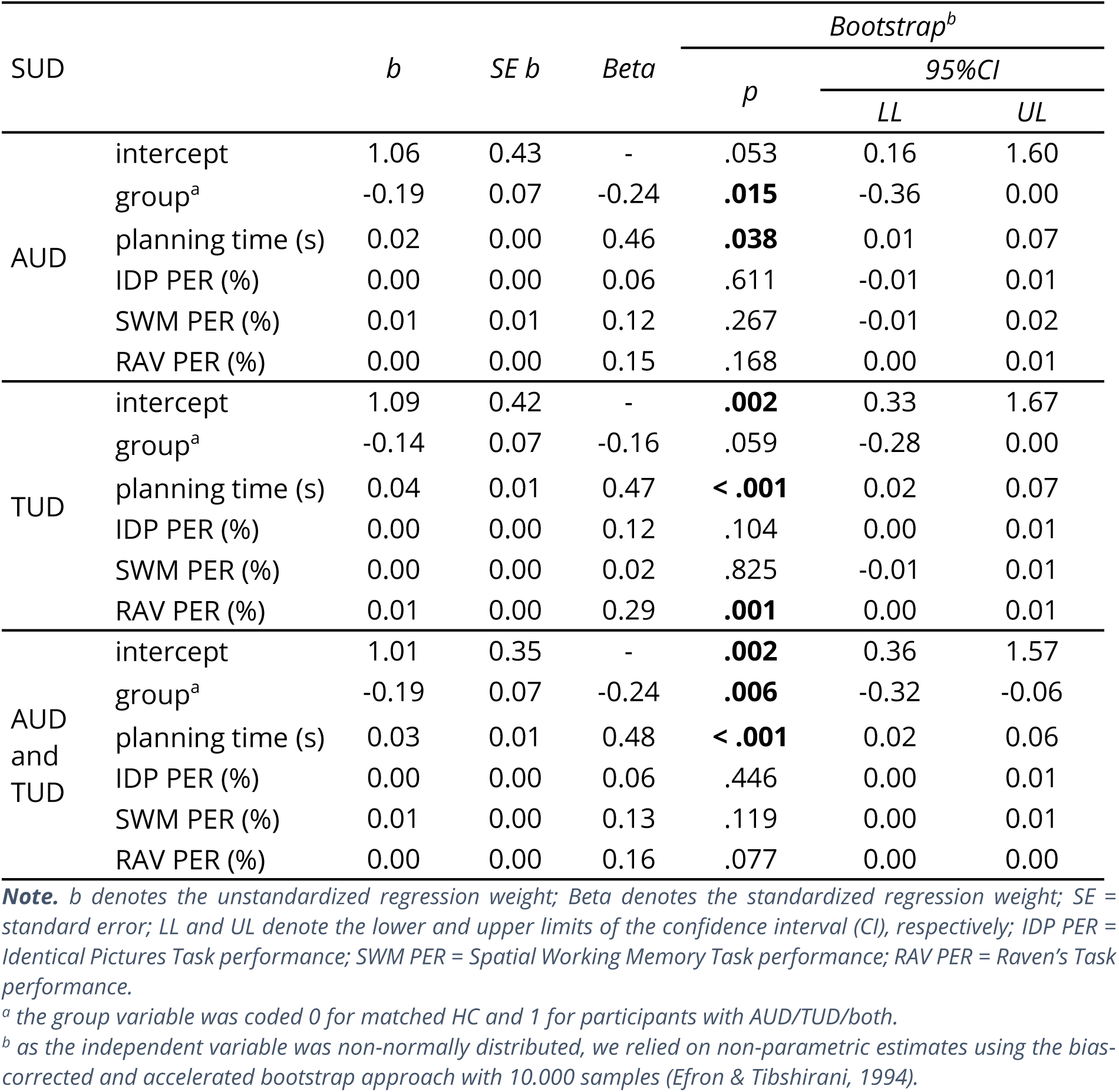
Estimates of Linear Regression Model for planning depth in the Space Adventure Task (SAT).

The factorial analysis revealed a significant compensatory interaction between AUD and TUD (b = 0.25, p = .044, BCa 95% CI [0.0, 0.50]), indicating that the combined effect of both disorders on planning depth was smaller than the sum of their individual effects. For planning time, SAT performance and the model parameters β and κ, there was no significant interaction effect. The analysis of the discounting parameter κ showed a significant effect of the TUD factor indicating that the group difference between HC and AUD and TUD is probably driven by the TUD condition. Details of the factorial analyses can be found in Table S3.

In terms of SUD severity and consumption levels, the post-hoc regression analyses (compare Table S4-S5) revealed a significant negative association of the sum of fulfilled TUD criteria with planning depth (*b* = -0.09, *p* = .047, BCa 95% CI [-.18, 0.0]). For the sum of fulfilled AUD criteria, there was a negative association with planning depth which was borderline significant (*b* = -0.07, *p* = .085, BCa 95% CI [-.14, 0.1]). Conversely, there was no significant association of planning depth with the number of heavy episodic drinking days or daily smoked cigarettes.

## DISCUSSION

In this study, we investigated forward planning in individuals with substance use disorders (SUD), focusing on the two most prevalent substances: alcohol use disorder (AUD) and tobacco use disorder (TUD). Across all groups, participants appeared to employ the same core heuristics previously identified in healthy individuals (Sass et al., 2025), including limiting planning depth, pruning low-probability branches, and probability discounting. This suggests that the fundamental cognitive strategies supporting resource-efficient planning are similar in individuals with SUD and healthy controls.

Despite this overall similarity, more detailed analyses revealed systematic differences in how these heuristics are applied. All three SUD groups consistently showed reduced planning depths relative to matched healthy controls (HC), indicating a shift in the extent of planning rather than in the type of strategies employed. Additionally, all SUD groups exhibited a tendency toward increased probability discounting, with the comorbid AUD+TUD group showing a significantly higher probability discounting compared to HC.

While earlier studies have already reported indications of incomplete planning in deterministic tasks with clinical AUD samples during or post-detoxification (Stephan et al., 2017), our findings extend this to a broader, general-population sample, and identify specific computational mechanisms of altered forward planning under uncertainty.

The association between SUD and reduced planning depth is a novel finding. Notably, our previous work found no reduction in planning depth among participants with predominantly mild-to-moderate AUD (Steffen et al., 2025). We attribute the present result to a larger sample size, greater severity of AUD cases, and improved methodological rigor through group matching. A possible interpretation of reduced planning depth in SUD is that it reflects impairments in working memory. However, performance on a spatial working memory task did not account for planning depth in our covariate analysis. Instead, the covariate analysis replicated the finding that planning depth is positively associated with planning time and logical reasoning across all participants. Shorter planning times may reflect reduced cognitive effort invested in the task, potentially through the evaluation of fewer segments of the decision tree. Additionally, the analysis showed that logical reasoning partially accounted for the reduced planning depth observed in the TUD group. This may reflect a selection bias toward participants with TUD and relatively lower fluid intelligence which would also explain why the TUD group selectively exhibited lower working memory and processing speed performance than matched healthy controls. More broadly however, we speculate that stronger logical reasoning abilities could facilitate more efficient forward planning by enabling participants to identify and prune branches of the decision tree that cannot yield better outcomes than those already evaluated. Nonetheless, the precise computational mechanisms linking reasoning ability and forward planning remain to be elucidated.

Our findings suggest that neurotoxic effects of alcohol or tobacco use may not fully explain the reduced planning depth observed in individuals with SUD. Although chronic alcohol and tobacco use have been shown to affect a broad range of cognitive functions (Durazzo et al., 2007, 2010; Stavro et al., 2013), individuals in our sample were relatively young (95% ≤ 45 years), well-functioning, and non–treatment seeking, a profile in which marked neurotoxic impairments are less common (Pfefferbaum et al., 1997). Moreover, we did not observe generalized impairments, and neither spatial working memory nor processing speed explained group differences in planning depth. The absence of additive reductions in planning depth in the comorbid AUD+TUD group, despite comorbidity often being associated with greater cognitive impairment (Durazzo et al., 2007; Hagger-Johnson et al., 2013; Pennington et al., 2013), further challenges a straightforward dose-dependent neurotoxicity account. Moreover, effect sizes for reduced planning depth were comparable across SUD groups, even though only the TUD group showed broader cognitive reductions. Such a pattern is difficult to reconcile with pure neurotoxicity and instead is consistent with the possibility that reduced planning depth may also reflect a pre-existing cognitive vulnerability to SUD more general. Supporting this interpretation, planning depth was more strongly associated with the number of fulfilled SUD criteria than with consumption levels. Because SUD criteria reflect maladaptive use rather than mere exposure, and prior work suggests only weak associations between SUD criteria and indices of heavy consumption (Oliver & Foulds, 2021; Tuithof et al., 2014), the association with planning depth may relate more to vulnerability toward SUD severity than to consumption-driven neurotoxicity. Nevertheless, longitudinal studies are required to test this hypothesis and to disentangle predisposing traits from consequences of prolonged substance use.

The fact that all groups, including those with SUD, adopted heuristics such as planning depth limitation and low-probability pruning supports the view that these strategies represent resource-rational approaches to complex planning (Sass et al., 2025). Following this reasoning, one might argue that minimal planning depth is in fact optimal, as simulations have shown that shallow planning strategies often yield the best cost-benefit ratio in this task (Sass et al., 2025). However, participants were explicitly instructed to plan ahead and maximize their points. Therefore, we interpret the planning depth reduction observed in SUD as an impairment in the ability to meet the task demands rather than an optimal strategy selection.

The third component of our planning model, probability discounting, reflects an undervaluation of probabilistic outcomes. In the context of our task and model assumptions, this component cannot be considered resource-rational, as it does not save any resources. All SUD groups showed a tendency toward stronger probability discounting, with the comorbid AUD+TUD group showing a significant effect. Previous work mainly focused on non-sequential decisions in SUD and found mixed results for AUD but more consistent evidence of increased risk aversion for probabilistic gains in TUD and SUD in general (for an overview see Garami & Moustafa, 2020; Harrison et al., 2018), which aligns with our finding in the domain of sequential planning. However, these studies often focused solely on decisions involving gains and did not consider probabilistic losses, which are known to elicit opposite behaviors (Kahneman & Tversky, 1979). In our task, probability discounting reflects a bias toward more certain gains and, simultaneously, an aversion to certain losses. This pattern suggests a lower sensitivity to probabilistic consequences in SUD which could underlie a stronger preference for substances of abuse which typically deliver reliable positive effects (e.g., euphoria, relief from distress), whereas negative consequences are less immediate, rarer, and harder to predict. Future research could explore this hypothesis by designing tasks that pit certain small rewards with large probabilistic losses against certain small losses with large probabilistic gains.

## Conclusion

In summary, our findings suggest that individuals with alcohol and/or tobacco use disorder engage in forward planning using generally similar strategies as healthy controls but show consistently reduced planning depth and a tendency toward stronger probability discounting. While the present results cannot determine whether these differences arise from neurotoxic effects, pre-existing traits, or both, the overall pattern is compatible with the possibility of a planning-related vulnerability that may contribute to SUD severity. This potential vulnerability offers a promising direction for future research and intervention.

## Limitations

A key limitation of this study was the lack of experimental control in the online setting, as participants completed the task at home. Although we implemented safeguards, such as instructing participants to perform the task in a quiet environment and to abstain from alcohol or other substances with long-term cognitive effects the day prior, we could not verify compliance or prevent distractions. More generally, we were unable to control for potential acute effects of recent substance use or withdrawal states, either of which could have influenced task performance. We attempted to exclude participants exhibiting random behavior, but future replications in controlled laboratory settings are necessary to rule out such confounds conclusively.

## AUTHOR CONTRIBUTION STATEMENT

All authors conceptualized the study design. M.N.S., S.S. and S.K. supervised the study execution and data analysis. J.S. implemented the study, conducted the assessments, performed computational modelling and data analysis, wrote the manuscript and designed the plots. All authors reviewed the manuscript.

## DATA AVAILABILITY

All data generated or analyzed during this study along with the code and scripts necessary to perform the model-based inference and statistical analyses are available in the plandepth_aud_tud Github repository, https://github.com/jeffensen/plandepth_aud_tud.

