## Supplementary Material for "Reduced planning depth and stronger probability discounting in alcohol and tobacco use disorder"

Michael N. Smolka

Full postal address:
Section of Systems Neuroscience
Department of Psychiatry
Technische Universität Dresden
Würzburger Str. 35
01187 Dresden (Germany)

**Funding**

This research was funded by the German Research Foundation (Deutsche Forschungsgemeinschaft, DFG project numbers 178833530 (SFB 940), 402170461 (TRR 265), 454245598 (IRTG 2773), and 521379614 (TRR 393)).

### ABSTRACT

***Table 1. Sample Characteristics of the three SUD groups and matched healthy controls.***

| | Alcohol | | | | Tobacco | | | | Both | | | |
|---|---|---|---|---|---|---|---|---|---|---|---|---|
| | AUD | HC | *t* | *p* | TUD | HC | *t* | *p* | AUD and TUD | HC | *t* | *p* |
| *N* | 45 | 45 | | | 49 | 49 | | | 45 | 45 | | |
| ***Sociodemography (matched)*** | | | | | | | | | | | | |
| Age | 30.2 (9.7) | 30.0 (10.3) | -0.10[a] | .923 | 31.0 (10.3) | 30.6 (10.0) | -0.35[a] | .725 | 28.1 (7.3) | 27.9 (7.3) | -0.15[a] | .878 |
| Gender (Male/Fem./Div.) | 23/22/- | 27/18/- | 0.72[b] | .396 | 29/20/- | 31/18/- | 0.17[b] | .678 | 27/17/1 | 26/19/0 | 1.13[b] | .568 |
| HEEQ (Yes/No) | 41/4 | 42/3 | 0.16[b] | .694 | 43/6 | 46/3 | 1.10[b] | .294 | 40/5 | 42/3 | 0.55[b] | .459 |
| Employed (Yes/No) | 35/10 | 36/9 | 0.67[b] | .796 | 35/14 | 34/15 | 0.05[b] | .825 | 38/7 | 40/5 | 0.39[b] | .535 |
| ***Psychological Measures*** | | | | | | | | | | | | |
| NFC score | 13.0 (15.9) | 15.9 (14.0) | 0.92 | .362 | 14.4 (14.4) | 12.8 (14.8) | 0.50[a] | .619 | 14.0 (12.3) | 11.7 (15.3) | -0.78 | .439 |
| Video games use | 2.5 (1.5) | 2.3 (1.4) | -0.49[a] | .622 | 2.6 (1.5) | 2.2 (1.6) | -1.2[a] | .230 | 2.7 (1.5) | 2.5 (1.3) | -0.35[a] | .725 |
| Electr. device use | 5.0 (0.0) | 5.0 (0.0) | 0.0[a] | 1.0 | 5.0 (0.1) | 5.0 (0.0) | -1.0[a] | .317 | 5.0 (0.0) | 5.0 (0.0) | 0.0[a] | 1.0 |
| ***Substance Use*** | | | | | | | | | | | | |
| AUD severity[d] | 0/0/12/33 | 45/0/0/0 | - | - | 49/0/0/0 | 49/0/0/0 | - | - | 0/10/18/17 | 45/0/0/0 | - | - |
| TUD severity[d] | 45/0/0/0 | 45/0/0/0 | - | - | 0/25/18/6 | 49/0/0/0 | - | - | 0/20/14/11 | 45/0/0/0 | - | - |
| AUDIT score | 18.0 (5.5) | 4.2 (3.5) | -8.02[a] | **<.001** | 8.3 (4.8) | 3.6 (3.8) | -5.28[a] | **<.001** | 16.3 (5.9) | 4.4 (3.5) | -7.59[a] | **<.001** |
| FTND score | 0.0 (0.3) | 0.0 (0.0) | -1.00[a] | .317 | 3.3 (2.2) | 0.0 (0.1) | -7.88[a] | **<.001** | 2.9 (2.3) | 0.0 (0.0) | -7.54[a] | **<.001** |
| HED days during last 3 months | 14.6 (13.8) | 1.8 (3.7) | -6.99[a] | **<.001** | 4.5 (5.5) | 1.1 (2.8) | -4.96[a] | **<.001** | 13.9 (15.6) | 1.9 (3.5) | -6.05[a] | **<.001** |
| Daily cigarettes last 3 months[c] | 1.8 (4.4) | 0.1 (0.3) | -3.61[a] | **<.001** | 12.5 (6.3) | 0.1 (0.4) | -8.83[a] | **<.001** | 14.1 (8.4) | 0.1 (0.3) | -8.55[a] | **<.001** |

***Note.*** *Descriptive statistics are formatted as: Mean (Standard Deviation).*

*HEEQ = German Higher Education Entrance Qualification; AUDIT = Alcohol Use Disorders Identification Test; HED = heavy episodic drinking; NFC = need for cognition.*

[a] *Z-statistic and corresponding asymptotic p-value: because of non-normality, a non-parametric Mann-Whitney-U test was performed.*

[b] *Pearson's $\chi^2$-statistic and corresponding p-value.*

[c] *Data missing for one participant of the AUD and TUD group and one participant of the AUD group due to technical failure.*

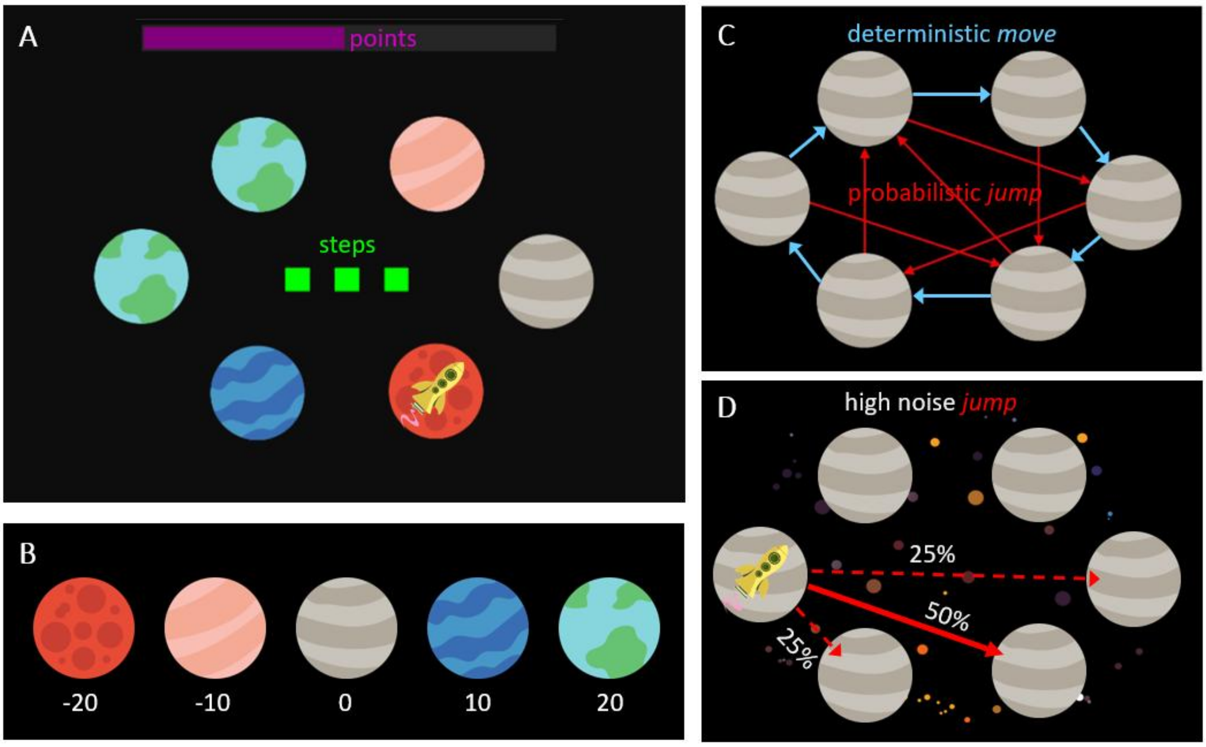


***Figure 1**. Schematic of the Space Adventure Task (adapted from Steffen et al., 2023).* ***(A)*** *Example mini-block with three action steps (green squares) under low-noise conditions (black background). The yellow rocket indicates the current location while the fuel bar at the top depicts total points accumulated so far throughout the task.* ***(B)*** *The five planet types with their respective reward values.* ***(C)*** *Transition rules for move (clockwise blue arrows) and*

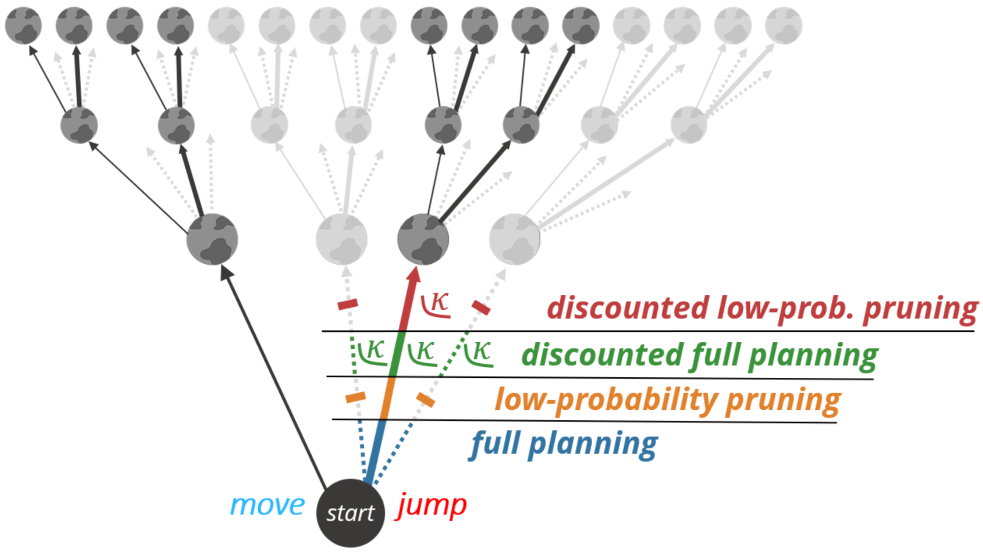


***Figure 2.*** *Visualization of the four forward planning strategies in the decision tree of the SAT. In each mini-block, participants could plan 3 steps ahead and choose between a deterministic move and a probabilistic jump action. The jump action led to the main target with a probability of p=0.9 or 0.5 (thick arrow) and to neighboring planets with a probability of p=0.05 or 0.25 each (dotted arrows). The full planning strategy (blue) evaluated all the branches of the tree with their accurate probabilities. The low-probability pruning strategy (orange) disregarded the low-probability transitions to the neighboring planets in all the levels of the decision tree (greyed out) and treated the transition to the main target as deterministic. The discounted full planning strategy (green) evaluated all branches but applied hyperbolic probability discounting to the jump transitions. Discounting was applied globally to the overall Q-value of the jump action with a participant-specific discounting parameter (κ). This corresponds to discounting the jump transitions of the first step in the decision tree. The discounted low-probability pruning strategy (red) combined probability discounting with pruning of the low-probability transitions. Thus, discounting was effectively only applied to the main transition of the first step in the decision tree with a participant-specific discounting parameter (κ).*

$$p(a_t = 'jump'|\, s_t, d) = \sigma(\beta \, * \, \Delta Q(s_t, d) + \theta), \tag{1}$$

where

$$\sigma(x) = \frac{1}{1 + e^{-x}} \tag{2}$$

$$\Delta Q(s_t, d) = Q(a_t = 'jump', s_t, d) - Q(a_t = 'move', s_t, d). \tag{3}$$

Here, we included two established parameters that further modify choice probabilities. First, the inverse response noise $\beta$ which controlled the extent to which differences in $Q$-values affected action selection. If $\beta = 0$, actions are selected with a constant probability independent of outcomes, while higher values of $\beta$ represented higher probability to select the action with the highest $Q$-value. Second, the parameter $\theta$ which denoted an a priori response bias, where positive values imply a bias towards choosing 'jump'. Hence, the full planning model contained three free parameters: $\beta$, $\theta$ and $d$.

**Discounted full planning**: this model extends the full planning model by probability discounting, i.e. a discounting of all outcomes of the probabilistic jump action. Discounting was applied at the root of the decision tree (see Figure 2), so all action sequences starting with a jump were discounted according to the probability of the jump transition. Therefore, this model differs from the full planning model in the computation of the relative $Q$-values:

$$\Delta Q(s_t, d) = \gamma_{disc} Q(a_t = 'jump', s_t, d) - Q(a_t = 'move', s_t, d) \tag{4}$$

Here, the expected cumulative reward of the jump action is discounted by a discounting factor $\gamma_{disc}$ which follows a typical hyperbolic discounting function (Green & Myerson, 2004):

$$\gamma_{disc} = \frac{p_{jump}}{p_{jump} + \kappa - \kappa p_{jump}} \tag{5}$$

where $p_{jump}$ denotes the probability of the high-probability jump transition $p(s_{t+1} = 'target'|s_t, a_t = 'jump')$ and $\kappa$ denotes the individual discounting parameter. In the case of $\kappa = 0$, there was no discounting. Higher values of $\kappa$ led to stronger probability discounting and, hence, to lower subjective values of all outcomes of the probabilistic jump action. The $\kappa$ -values were limited at a maximum of 30, as values beyond this threshold did not offer additional information. The discounted full planning model contained four free parameters: $\beta$, $\theta$, $\kappa$ and $d$.

*Table 2. Descriptive Statistics and group comparison of task outcomes and model parameters.*

| | Alcohol | | | | Tobacco | | | | Both | | | |
|---|---|---|---|---|---|---|---|---|---|---|---|---|
| | AUD | HC | *t* | *p* | TUD | HC | *t* | *p* | AUD and TUD | HC | *t* | *p* |
| *N* | 45 | 45 | | | 49 | 49 | | | 45 | 45 | | |
| **Task Performances [%]** | | | | | | | | | | | | |
| Space Adventure | 67.1 (14.5) | 69.9 (10.1) | -0.55[a] | **.002** | 67.0 (8.8) | 70.2 (10.6) | -2.31[a] | **.021** | 66.2 (11.4) | 70.9 (9.5) | -2.15[a] | **.032** |
| Identical Pictures | 75.7 (10.2) | 74.7 (12.9) | -0.41 | .680 | 70.0 (11.1) | 75.3 (12.0) | -2.57[a] | **.010** | 74.5 (10.1) | 75.6 (12.5) | -0.91[a] | .363 |
| Spatial Working Memory | 89.1 (7.5) | 89.0 (7.4) | -0.12[a] | .907 | 86.8 (7.2) | 88.5 (9.0) | -2.32[a] | **.020** | 87.3 (8.4) | 88.3 (8.3) | -0.91[a] | .365 |
| Raven's Matrices | 45.0 (20.2) | 57.2 (22.4) | 2.72 | **.008** | 39.3 (23.1) | 56.5 (23.2) | -3.70[a] | **<.001** | 43.0 (25.8) | 59.3 (21.8) | -3.01[a] | **.003** |
| **Task Reaction times [s]** | | | | | | | | | | | | |
| Space Adventure[b] | 7.6 (9.5) | 8.6 (5.7) | -1.68[a] | **.019** | 7.5 (4.7) | 8.3 (5.3) | -0.85[a] | .396 | 7.0 (5.2) | 8.7 (5.3) | -1.97[a] | **.048** |
| Identical Pictures | 2.3 (.4) | 2.3 (.5) | -0.03[a] | .974 | 2.5 (0.4) | 2.3 (0.5) | -2.58[a] | **.010** | 2.2 (0.3) | 2.3 (0.5) | -0.37[a] | .713 |
| Spatial Working Memory | 1.3 (.3) | 1.3 (.4) | -1.32[a] | .187 | 1.4 (0.3) | 1.3 (0.3) | -2.50[a] | **.012** | 1.3 (0.3) | 1.3 (0.3) | -1.12[a] | .264 |
| **Model Parameters** | | | | | | | | | | | | |
| Planning Depth | 1.9 (.4) | 2.2 (.4) | 3.0[a] | **.003** | 1.9 (0.5) | 2.1 (0.3) | -3.03[a] | **.002** | 1.9 (0.4) | 2.2 (0.3) | -3.35[a] | **.001** |
| Inv. Resp. Noise $\beta$ | 1.8 (.6) | 1.9 (.4) | -1.23[a] | .218 | 1.8 (0.4) | 1.9 (0.4) | 1.21 | .229 | 1.7 (0.4) | 1.9 (0.3) | 2.62 | **.010** |
| Response Bias $\theta$ | -0.2 (.3) | -0.3 (.3) | -1.69 | .095 | -0.2 (0.3) | -0.2 (0.3) | 0.08 | .938 | -0.2 (0.3) | -0.3 (0.3) | -0.91 | .366 |
| Discounting $\kappa$ | 4.1 (4.5) | 3.0 (3.6) | -1.70[a] | .089 | 5.0 (5.3) | 2.7 (3.2) | -1.62[a] | .104 | 6.6 (7.0) | 2.6 (2.9) | -2.28[a] | **.023** |

***Note.*** *Descriptive statistics are formatted as: Mean (Standard Deviation).*

[a] *Z-statistic and corresponding asymptotic p-value: because of non-normality, a non-parametric Mann-Whitney-U test was performed.*

[b] *i.e. planning time*

#### Model Comparison of Planning Strategies

The comparison of the four different planning strategy models revealed that the combined model with pruning of low-probability branches as well as probability discounting fit the data best (see Figure 3). All further analyses of model parameters are therefore based on this winning model. Separate model comparison for each SUD group and the pooled group of all HC participants included through matching showed similar results for each group (see Figure S4). Only for the AUD group, the fit of the discounted low-probability pruning model was indistinguishable from the full planning and discounted full planning model, so there was no single best fitting model.

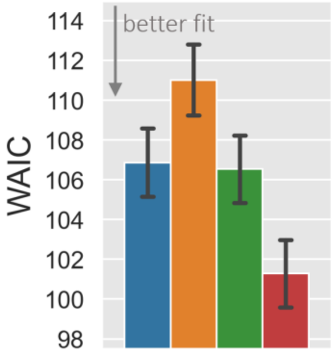


| | $WAIC_{model}$ | | $\Delta WAIC_{fp-model}$ | |
|---|---|---|---|---|
| **Model** | Mean | *SE* | Mean | *SE* |
| full planning (fp) | 105.70 | 1.29 | - | - |
| low-probability pruning | 108.87 | 1.35 | -4.16 | 1.07 |
| discounted full planning | 105.37 | 1.26 | 0.33 | 0.51 |
| discounted low-prob. pruning | 99.70 | 1.27 | 5.59 | 1.01 |

***Figure 3***. *Model comparison of the four planning strategies with the Widely Applicable Information Criterion (WAIC), across SUD participants and HC participants included during matching. The full planning strategy (fp, blue) evaluated all the branches of the decision tree with their accurate probabilities. The low-probability pruning strategy (orange) disregarded the low-probability transitions to the neighboring planets. The discounted full planning strategy (green) evaluated all branches but applied hyperbolic probability discounting to the jump transitions and the discounted low-probability pruning strategy (red) combined probability discounting with pruning of the low-probability transitions. Error bars indicate the standard error of the mean of participants' values (SE). The barplot's values are also depicted in the $WAIC_{model}$ column table. The last column of the table presents the mean pairwise WAIC differences of the participants' values between the null model (full planning) and each of the other models alongside the corresponding standard error of the mean (SE).*

#### Planning Depth and model parameters

The distribution and group comparison of all model parameters is illustrated in Figure 4. The planning depth parameter was reduced in all subgroups with SUD compared to their matched HC group, i.e. for AUD ($Z$ = 3.00, $p$ = .003), TUD ($Z$ = -3.03, $p$ = .002) as well as AUD and TUD ($Z$ = -3.35, $p$ = .001). Post-hoc pairwise Z-tests revealed that there was no significant difference between any of these three effects (see Table S2). Furthermore, we found that the comorbid group with AUD and TUD also showed higher response noise as indicated by the parameter $\beta$ ($Z$ = 2.62, $p$ = .010) and stronger probability discounting as indicated by the discounting parameter $\kappa$ ($Z$ = -2.28, $p$ = .023) as the HC participants. Across all participants, there was a strong positive correlation of performance in the planning task with planning depth ($r$ = 0.63, $p$ < .001) as well as with inverse response noise $\beta$ ($r$ = 0.72, $p$ < .001) and a strong negative correlation with the discounting parameter $\kappa$ ($r$ = -0.60, $p$ < .001).

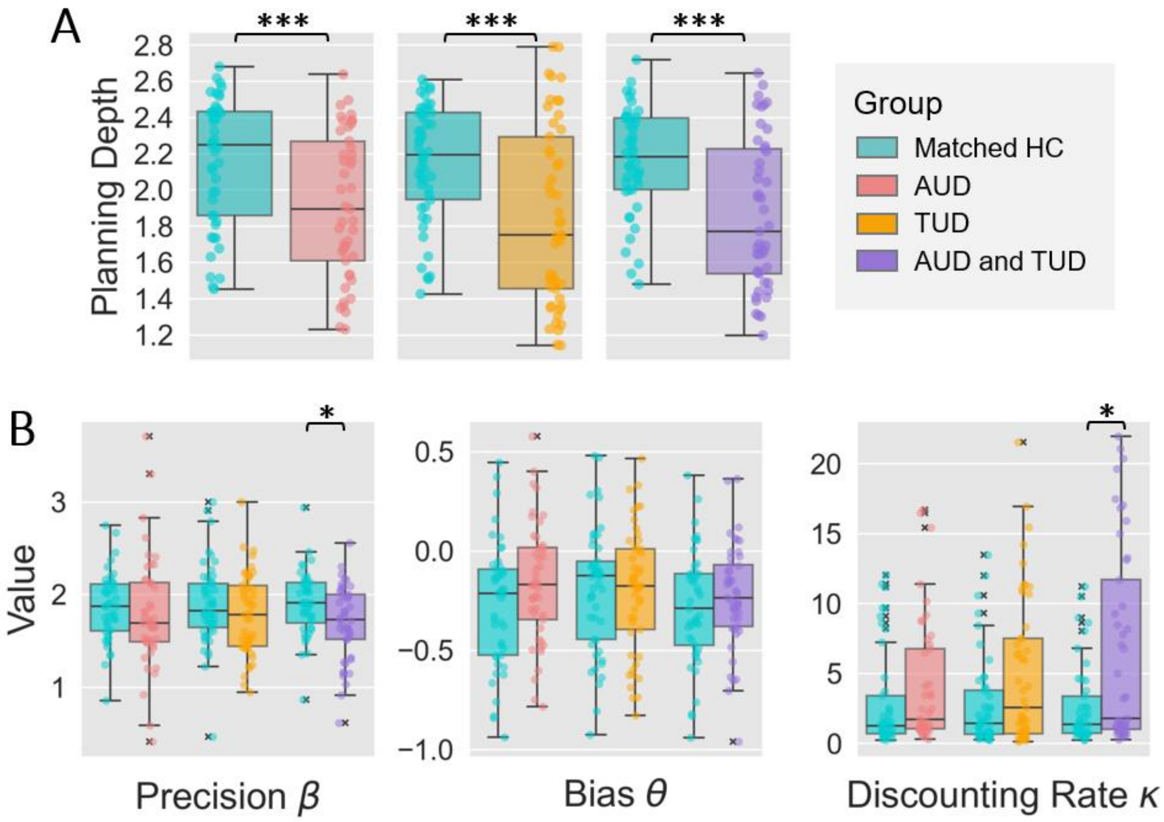


***Figure 4.*** *Boxplots of inferred model parameter values per subject comparing the groups with AUD/TUD/both to their respective matched healthy control (HC) group.* ***(A)*** *Averaged planning depths per subject.* ***(B)*** *Additional model parameters precision β, bias θ and probability discounting rate κ per subject.*

To analyze the effect of the noise condition on planning depth, we performed separate linear mixed effects model analyses for each SUD group also showing reduced planning depths in SUD compared to matched control participants. Moreover, the analyses revealed that across groups participants, under high noise, marginally reduced their planning depth on average by 0.06. This noise effect as well as overall mean of planning depths represented by the intercept showed considerable variation among participants as indicated by the variance parameter $\eta$. For detailed statistics, please refer to Table 3.

***Table 3. Estimates of Linear Mixed Effects Model for Planning Depth***

| SUD | | *b* | *SE b* | *p* | *η* | *SE η* | *p* |
|---|---|---|---|---|---|---|---|
| AUD | intercept | 2.18 | 0.06 | **< .001** | 0.14 | 0.02 | **< .001** |
| | group[a] | -0.24 | 0.08 | **.003** | - | - | - |
| | noise | -0.06 | 0.01 | **< .001** | 0.00 | 0.00 | .083 |
| | group x noise | 0.00 | 0.01 | .991 | - | - | - |
| TUD | intercept | 2.17 | 0.06 | **< .001** | 0.17 | 0.02 | **< .001** |
| | group[a] | -0.29 | 0.08 | **< .001** | - | - | - |
| | noise | -0.06 | 0.01 | **< .001** | 0.00 | 0.00 | **< .001** |
| | group x noise | 0.01 | 0.01 | .342 | - | - | - |
| AUD and TUD | intercept | 2.19 | 0.05 | **< .001** | 0.13 | 0.02 | **< .001** |
| | group[a] | -0.30 | 0.08 | **< .001** | - | - | - |
| | noise | -0.05 | 0.01 | **< .001** | 0.00 | 0.00 | **.028** |
| | group x noise | 0.01 | 0.01 | .401 | - | - | - |

***Note.*** *b denotes the fixed effect parameter of the statistical model; η denotes the random effect parameter; SE = standard error.*
*[a] the group variable was coded 0 for matched HC and 1 for participants with AUD/TUD/both.*

To investigate the relationship of planning depth with planning time and the cognitive tasks, we performed a multiple linear regression for each SUD group and relied on non-parametric bootstrapping (BCa; Efron & Tibshirani, 1994) because of non-normal distributions of planning depths (see Table 4). For all SUD groups, we found a significant positive association of planning time and planning depth, i.e. for the AUD ($p$ = .038, BCa 95% CI [0.01, 0.07]), TUD ($p$ < .001, BCa 95% CI [0.02, 0.07]) and the AUD and TUD group ($p$ < .001, BCa 95% CI [0.02, 0.06]). Despite the inclusion of the additional predictors, the group effect of planning depth was still significant for the AUD group ($p$ = .015, BCa 95% CI [-0.36, 0.0]) and the AUD and TUD group ($p$ = .006, BCa 95% CI [-0.32, -0.06]). In the TUD group, we found a significant positive association with the logical reasoning performance in the raven's task ($p$ = .001, BCa 95% CI [0.0, 0.01]) and here, the group effect got borderline significant when accounting for planning time and performances in cognitive tasks ($p$ = .059, BCa 95% CI [-0.28, 0.0]). Across all participants, there were positive correlations of planning depth with planning time ($r$ = 0.52, $p$ < .001) as well as with logical reasoning ($r$ = 0.38, $p$ < .001; see FIGURE S5 for a full correlation table).

***Table 4. Estimates of Linear Regression Model for planning depth in the Space Adventure Task (SAT).***

| SUD | | *b* | *SE b* | *Beta* | Bootstrap[b] | | |
|---|---|---|---|---|---|---|---|
| | | | | | *p* | *95%CI* | |
| | | | | | | *LL* | *UL* |
| AUD | intercept | 1.06 | 0.43 | - | .053 | 0.16 | 1.60 |
| | group[a] | -0.19 | 0.07 | -0.24 | **.015** | -0.36 | 0.00 |
| | planning time (s) | 0.02 | 0.00 | 0.46 | **.038** | 0.01 | 0.07 |
| | IDP PER (%) | 0.00 | 0.00 | 0.06 | .611 | -0.01 | 0.01 |
| | SWM PER (%) | 0.01 | 0.01 | 0.12 | .267 | -0.01 | 0.02 |
| | RAV PER (%) | 0.00 | 0.00 | 0.15 | .168 | 0.00 | 0.01 |
| TUD | intercept | 1.09 | 0.42 | - | **.002** | 0.33 | 1.67 |
| | group[a] | -0.14 | 0.07 | -0.16 | .059 | -0.28 | 0.00 |
| | planning time (s) | 0.04 | 0.01 | 0.47 | **< .001** | 0.02 | 0.07 |
| | IDP PER (%) | 0.00 | 0.00 | 0.12 | .104 | 0.00 | 0.01 |
| | SWM PER (%) | 0.00 | 0.00 | 0.02 | .825 | -0.01 | 0.01 |
| | RAV PER (%) | 0.01 | 0.00 | 0.29 | **.001** | 0.00 | 0.01 |
| AUD and TUD | intercept | 1.01 | 0.35 | - | **.002** | 0.36 | 1.57 |
| | group[a] | -0.19 | 0.07 | -0.24 | **.006** | -0.32 | -0.06 |
| | planning time (s) | 0.03 | 0.01 | 0.48 | **< .001** | 0.02 | 0.06 |
| | IDP PER (%) | 0.00 | 0.00 | 0.06 | .446 | 0.00 | 0.01 |
| | SWM PER (%) | 0.01 | 0.00 | 0.13 | .119 | 0.00 | 0.01 |
| | RAV PER (%) | 0.00 | 0.00 | 0.16 | .077 | 0.00 | 0.00 |

***Note.*** *b denotes the unstandardized regression weight; Beta denotes the standardized regression weight; SE = standard error; LL and UL denote the lower and upper limits of the confidence interval (CI), respectively; IDP PER = Identical Pictures Task performance; SWM PER = Spatial Working Memory Task performance; RAV PER = Raven's Task performance.*

*[a] the group variable was coded 0 for matched HC and 1 for participants with AUD/TUD/both.*

*[b] as the independent variable was non-normally distributed, we relied on non-parametric estimates using the bias-corrected and accelerated bootstrap approach with 10.000 samples (Efron & Tibshirani, 1994).*

#### COMPETING INTERESTS

The authors declare no competing interests.

#### DATA AVAILABILITY

All data generated or analyzed during this study along with the code and scripts necessary to perform the model-based inference and statistical analyses are available in the plandepth_aud_tud Github repository, https://github.com/jeffensen/plandepth_aud_tud.
